# Associated habitat and suitability modeling of goldenseal (*Hydrastis canadensis* L.) in Pennsylvania: explaining and predicting species distribution in a northern edge of range state

**DOI:** 10.1101/694802

**Authors:** Grady H. Zuiderveen, Xin Chen, Eric P. Burkhart, Douglas A. Miller

## Abstract

Goldenseal (*Hydrastis canadensis* L.) is a well-known perennial herb indigenous to forested areas in eastern North America. Owing to conservation concerns including wild harvesting for medicinal markets, habitat loss and degradation, and an overall patchy and often inexplicable absence in many regions, there is a need to better understand habitat factors that help determine the presence and distribution of goldenseal populations. In this study, flora and edaphic factors associated with goldenseal populations throughout Pennsylvania—a state near the northern edge of its range—were documented and analyzed to identify habitat indicators and provide possible in situ stewardship and farming (especially forest-based farming) guidance. Additionally, maximum entropy (Maxent) modeling was applied to better predict where suitable habitat might be encountered more broadly and explain species absence from regions of the state (and the northeastern states). Habitat study results identified rich, mesic, woodland sites as being suitable for goldenseal. The most prevalent overstory tree associates on such sites were tulip-poplar (*Liriodendron tulipifera* L.) and sugar maple (*Acer saccharum* Marshall), and the most common understory associates were spicebush (*Lindera benzoin* L.), Virginia creeper (*Parthenocissus quinquefolia* (L.) Planch.), Jack-in-the-pulpit (*Arisaema triphyllum* (L.) Schott), mayapple (*Podophyllum peltatum* L.), wood fern (*Drypoteris marginalis* (L.) A. Gray), and rattlesnake fern (*Botrypus virginianus* (L.) Michx.). Loam soils were the most common textural class (average sand, silt, and clay ratio of 50:30:20) with an average pH of 6.2 and high variation in macronutrients. While such sites are widespread in the state, Maxent modeling suggested the present distribution in Pennsylvania is largely restricted by winter temperatures and bedrock type. The latter of these, in turn, is correlated in part with land use legacy (e.g., clearing for farming or livestock grazing), especially in southeast portions of the state.

## Introduction

The Appalachian Mountains of eastern North America, which span from Alabama, United States to New Brunswick, Canada, are a botanical diversity hotspot due to variations in elevation, moisture, and latitude (Pickering et al., 2002). However, much of this biodiversity is threatened from wild plant harvesting, land development, site degradation and invasive species (Pickering et al., 2002; Cruse-Sanders and Hamrick, 2004; Wickham et al., 2007). One such species that is of conservation concern within this region is goldenseal (*Hydrastis canadensis* L.) (Oliver, 2017; NatureServe, 2018; Oliver and Leaman, 2018).

Goldenseal is a well-known medicinal herb indigenous to the forests of eastern North America, from southern Canada to northern Georgia. Populations can be found on a variety of different sites from open woods on hillsides and ridges, to lowland wooded sites along streams and rivers where there is good drainage (McGraw et al., 2003; Predny and Chamberlain, 2005; NatureServe, 2018). Reproducing goldenseal populations have been found growing on very different geologic substrates, ranging from a calcareous, dolomitic limestone site with a pH of 4.4 to 8.0 to a more mountainous granitic terrain with a pH range of 5.5 to 6.0 (Mueller, 2004). Some authors note that beyond the broad variation in reported sites, the species seems to prefer well-drained, moist, calcareous soils, high in organic matter, with pH ranging from 5.5-6.5 (Upton, 2001; Sinclair and Catling, 2001).

Goldenseal is inexplicably rare in many parts of its range, despite the availability of seemingly suitable habitat, and researchers have struggled to identify what factors determine or restrict its distribution across Appalachia (McGraw et al., 2003; Sanders, 2004; Sanders and McGraw, 2005). Goldenseal utilizes a mixed breeding system, which makes it unlikely that a lack of fruit production in isolated populations is contributing to rarity (Sanders, 2004). Additionally, the species can colonize habitats through colonial expansion through rhizome and root fragments (Van Fleet, 1914; Davis and McCoy, 2000). In recent decades, some authors and observers have attributed goldenseal’s rarity in many otherwise suitable forests to habitat destruction and exploitation by commercial harvesters (Mulligan and Gorchov, 2004; Sanders, 2004; Albrecht and McCarthy, 2006; Rhoads and Block, 2007).

In addition to explaining the patchy distribution of goldenseal, there is increased interest in growing goldenseal as a commercial crop, especially amongst forest landowners. Although goldenseal can be grown using field-based cultivation under artificial shade, cultivating the plant using forest-based farming approaches (i.e., agroforestry) has potential advantages over field based cultivation. These include cost savings associated with artificial shade structures (goldenseal is shade obligate) and possible reduction in disease, a higher incidence of which can be associated with field grown production (Sinclair and Catling, 1996; Burkhart and Jacobson, 2009). In a study of woodland owners in Pennsylvania, just over a third (36%) of respondents were interested in using agroforestry for specialty crop production, including forest farming of medicinal plants (Strong and Jacobson, 2006). In a national survey, 75% of respondents were interested in learning how to manage their property for non-timber forest products (NTFPs) (McLain and Jones, 2013). By assisting landowners to successfully adopt forest farming of plants such as goldenseal, market demand could be met with a more consistent and sustainable supply of raw materials, and thereby reduce pressure on wild populations by easing demand for wild harvested materials.

One important component of successful in situ conservation and forest-based cultivation is to develop state level population and habitat data that can inform landscape management decisions and site selection for forest farming. This guidance and modeling can assist in identifying populations and areas for preservation and conservation, and can be used to identify landscapes that may be suitable for reintroduction (Questad et al., 2014) and forest farming, without extensive field surveying (Raxworthy et al., 2003). However, due to the rarity of some plant species, including goldenseal, traditional presence/absence modeling is not well suited, and traditional stratified random sampling has limited application (McGraw et al., 2003).

Goldenseal presence data are limited due to land ownership access limitations and lack of botanical surveys across the state; similarly, absence data is complicated by a host of historical and contemporary influences that are difficult to account for. For example, the absence of goldenseal at a site could be the result of previous harvest activities or land use legacies (Bellemare et al., 2002) rather than any habitat influences. Utilizing a modeling approach that requires presence-only data can be more suitable. One commonly used method, Maxent (Phillips et al., 2006) has been found to outperform other presence-only, as well as presence/absence modeling techniques such as GLM (Hirzel and Guisan, 2002) GAM (Yee and Mitchell, 1991), and BIOCLIM (Busby, 1991), and is particularly effective when presence data is limited (Elith et al., 2006; Hernandez et al., 2006; Wisz et al., 2008; Razgour et al., 2011; Yi et al., 2016). Maxent is a program that utilizes the principle of maximum entropy to model species distributions by minimizing relative entropy between a set of presence data and the landscape given a set of predictor variables (Elith et al., 2011). The principle of maximum entropy states that the probability distribution that best represents the actual distribution is the one with the largest entropy (i.e. highest average rate of information produced by a random source of data).

Combining state level predictive modeling with a field-based approach to assess microsite habitat conditions could be a particularly useful strategy to help inform a variety of land managers, from forest planners to and private forest landowners. Specifically, utilizing “indicator species” (i.e., nearest neighbors and associates) for on the ground site assessments can guide rapid assessment of suitable habitat of species of conservation concern (Burkhart, 2013), since forest vegetation is in part a result of the underlying climatic, topographic, and edaphic factors (Gilliam, 2014). Indicator species are often inadequate predictors of site quality when used in isolation (Turner and McGraw, 2015), but in combination with habitat suitability modeling broader predictive guidance could be developed. If indicator species are identified that respond similarly to habitat cues and conditions, stakeholders could use these species, and presence only models based on these species, as identifiers for areas suitable for surveying and conservation or farming reintroductions (Ren et al., 2010).

The goal of the study was to combine indicator species analysis (McCune et al., 2002) with habitat suitability modelling that incorporates climatic, edaphic, and topographic factors to provide a more robust understanding of potential goldenseal habitat in the state, and more broadly the northern edge of range for this species. Maxent was used to develop a habitat suitability model for goldenseal in Pennsylvania (PA) as the state is well suited for the application of habitat suitability mapping as it is diverse geographically, and is near the northern edge of the species native range, which could limit the effectiveness of a range wide suitability map (Braunisch et al., 2008). Given that the species is listed as “PA vulnerable” (PA DCNR, 1982), results may aid in population and habitat conservation goals, and provide guidance for forest farming adoption in Pennsylvania and mid-Atlantic and northeastern region more broadly.

## Materials and Methods

### Study Area

This study focused on goldenseal habitat in Pennsylvania, U.S.A. (39°43′-42°16′ N; 74°41′-80°31′ W). The apparent transition within PA from numerous occurrences in the southern part of the state to no known occurrences in the northern third of the state provides a unique opportunity to determine the influence of climate on goldenseal habitat suitability. Further, the state varies greatly in physiographic features including the Allegheny plateau in the western part of Pennsylvania, the Ridge-and-Valley region of central Pennsylvania, and the Piedmont region in the eastern part of Pennsylvania, which provides opportunity for identifying important edaphic and topographic factors on goldenseal habitat suitability.

In 2015 and 2016, wild and forest farmed goldenseal sites were solicited from botanist, goldenseal collectors, and forest landowners in PA. Ultimately, 27 sites and 58 plots were identified for inclusion in the study. Two of the sites were forest farmed populations but were included as one of the main objectives of the study was to determine suitable habitat for forest-based goldenseal cultivation. The sites were located throughout the lower two thirds of the state within 19 of PA’s 67 counties, two of which goldenseal had not been previously reported. Ten sites were located on the Allegheny plateau, five sites were in the Ridge-and-Valley, and 12 sites were from the Piedmont region (Figure 1).

**Figure 5.1.**
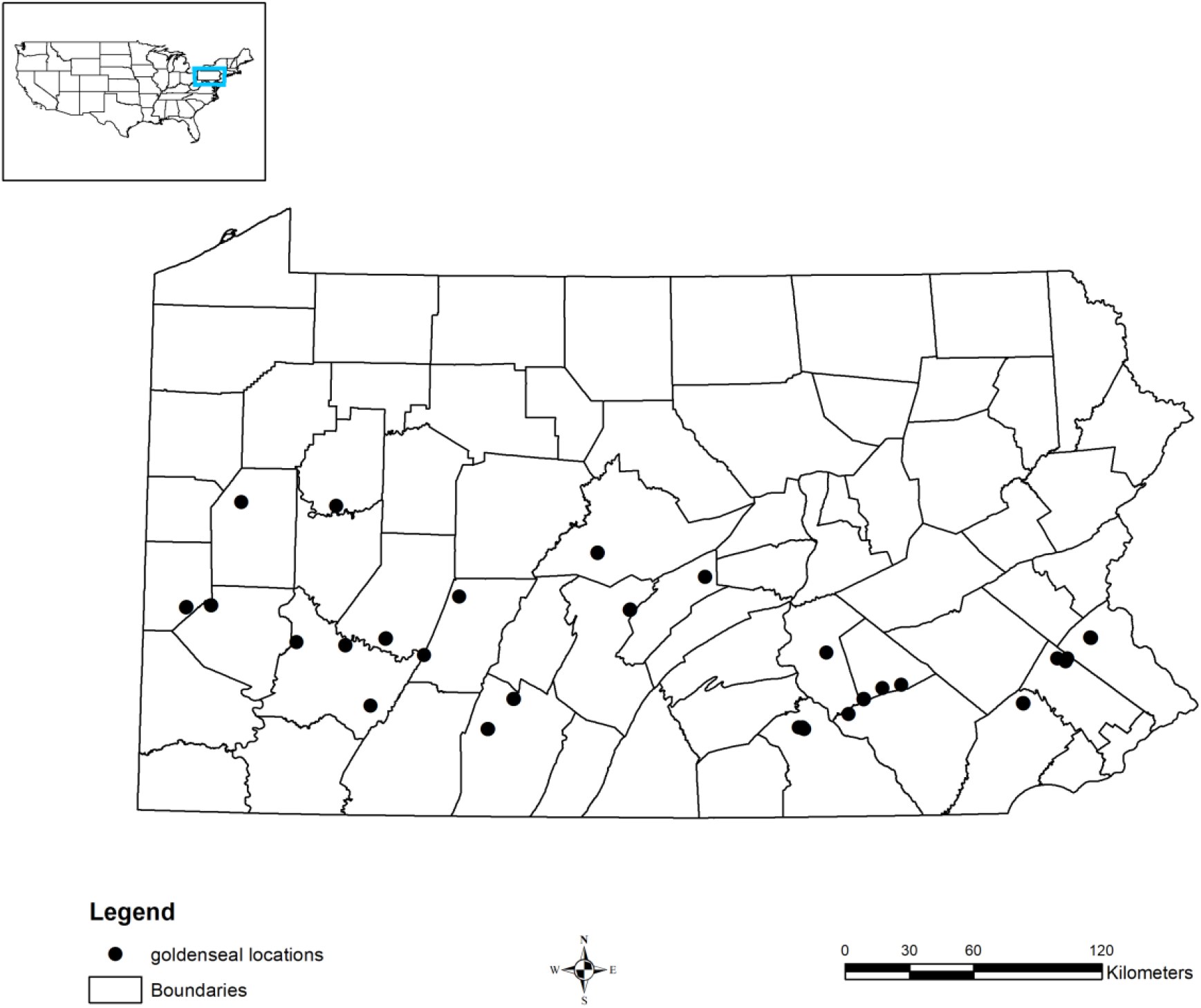
Distribution of goldenseal plots across Pennsylvania. All points are fuzzy as to not reveal the exact location of the goldenseal populations.

### Vegetation Sampling

Within each study site, plots were placed in as many as four discretely separated colonies. The number of plots varied due to overall population size and number of discrete colonies. Inclusion was restricted to reproducing populations. Some populations included in this study consisted of thousands of ramets spread over multiple hectares, while others consisted of a single colony of roughly 100 ramets.

Plots were established at each site centered on spatially discrete colonies of goldenseal, in a visually guided fashion intended to capture any differences in edaphic and floristic variables across each occurrence. The objective was to document only the vegetation nearest to, and interspersed with each goldenseal colony (i.e., “nearest neighbor”). To accomplish this, at each plot, overstory dominant and co-dominant tree species were recorded using the point-centered quarter method (PCQM). Using this method, the nearest tree species in each quarter was recorded for a total of four trees per plot. For each tree, distance from plot center and diameter at breast-height (DBH) was recorded for determination of importance values (Curtis and McIntosh, 1951; McCune et al., 2002).

Inventory of understory vegetation—including shrubs, ferns, and herbaceous species— was conducted using a fixed area circular plot with a radius of 5.64 meters (roughly 1/100 hectare). Each plot was visited at least twice during 2016 and 2017 to collect floristic data both in spring and summer to capture seasonal changes in associated flora.

Herbarium voucher specimens were collected for goldenseal from each plot and deposited at the Pennsylvania State University Herbarium (PAC), the Carnegie Museum of Natural History Herbarium (CM), and the Morris Arboretum of the University of Pennsylvania Herbarium (MOAR). Global positioning system (GPS) coordinates for all study plots are not on these vouchers to protect the exact location but area available from the Pennsylvania Department of Conservation and Natural Resources Wild Plant Management Program.

### Soil Analysis

Soil samples were collected from the upper 10 cm (A-horizon) after removing large coarse organic matter from each sample plot. Samples were sent to the Penn State Agricultural Analytical Services Lab for soil chemistry testing. Results include cation exchange capacity (CEC), water pH, phosphorus, potassium, magnesium, and calcium by the Mehlich 3 (ICP) test. Soil texture samples were collected from only a subset of the sites (n=19, 65% of sites).

### Statistical Analysis of Habitat Data

Descriptive statistics were generated for floristic and edaphic data from each plot. Indicator species analysis (ISA) (McCune et al., 2002) was used to determine if associated species were different across physiographic province, soil calcium content, and/or pH. Thresholds for edaphic factors were based on guidelines to maintain a mixed-species woodlot as provided by the Pennsylvania State Soil Analytics Lab. A Monte Carlo randomization procedure (with 4,999 randomizations) was used to determine significance (McCune et al., 2002). All ISA analysis was done using PC-ORD (Multivariate Analysis of Ecological Data, v. 6.0, MJM software design, Gleneden Beach, Oregon). Additionally, importance values were calculated on dominant or co-dominant tree species based on relative frequency, relative density, and relative dominance (Curtis and McIntosh, 1951).

### Modeling Habitat Suitability

Maximum entropy modeling (Maxent) was used to construct the habitat suitability model (Phillips et al., 2006) with presence-only data. Maxent version 3.4.1 was used to fit the model, using the default model parameters, which have been shown to perform well with a small sample size (Phillips and Dudík, 2008). In the model, given an unknown population, over a set number of background points, an approximate population probability distribution can be estimated from the sample probability distribution. In other words, the model minimizes the relative entropy between the probability estimated from the presence data and the probability estimated from the landscape defined in a space populated by the covariates, or independent variables (Elith et al., 2011). Specific to this study, a probability distribution for goldenseal habitat suitability was developed using the goldenseal presence data mapped against 10,000 randomly selected background points across Pennsylvania. The training data were comprised of 90% of the sample data randomly selected, and the test data were the remaining 10% of the sample data. The habitat suitability curves for each predictor variable were calculated and their contribution estimated using a jackknife test.

### Predictor Variables

In the Maxent model, 50 predictor variables were used to characterize abiotic conditions including soil, topography, and climate (Table B1). After preliminary analysis, the ten most influential variables were selected for the final model, to reduce overfitting due to a small sample size (Table 5.1). Topographic variables were computed from a 1 m digital elevation model available from Pennsylvania Spatial Data Access (http://www.pasda.psu.edu), while the soil moisture index was developed by Iverson et al. (1997). Edaphic features represented the top 30 cm of the soil and were derived from the Gridded SSURGO (gSSURGO) dataset (Soil Survey Staff 2017). Climatic predictors were obtained from WorldClim, and contained variables that represent annual trends, seasonality, and extreme environmental factors at 1 km spatial resolution. To map goldenseal habitat, the topographic predictors, integrated soil moisture index, and edaphic predictors were resampled to 90 m resolution using ESRI ArcGIS^©^.

**Table 5.1.** Predictor variables used to develop the habitat suitability model in Maxent.

| Variable | Description/Source |
| --- | --- |
| Climatic predictors |  |
| BIO4 | Temperature Seasonality (standard deviation *100) |
| BIO5 | Mean maximum temperature of the warmest month in degrees C |
| BIO11 | Mean temperature of coldest quarter degrees C |
| Edaphic predictors |  |
| BEDROCK | Bedrock type |
| SOILORD | Soil order based on USDA soil taxonomy |
| SOILSUBORD | Soil suborder based on USDA soil taxonomy |
| Topographic predictors |  |
| IMI | Integrated Moisture Index (Iverson et al., 1997) |
| ROUGH15 | Variance of elevation (15-pixel neighborhood) |
| ASPECT | Transformed aspect according to Beer et al. (1966) |
| ERODIBILITY | Soil erodibility factor |

## Results and Discussion

### Floristic Associations

A total of 159 species were documented in the study plots: 108 herbaceous plants; seven ferns; 20 vines, shrubs, and understory trees; and 24 overstory trees (Table A2). The most common herb was Jack-in-the-pulpit (*Arisaema triphyllum* (L.) Schott), which occurred in 79% of the plots. Only 13 additional herbaceous species occurred in more than 40% of the plots. Of the 108 herbs recorded, 72 of them occurred on less than 20% of the plots. Indicator species analysis identified physiographic province, soil calcium, and soil pH as significant. Of the 36 herbs that were present in more than 20% of the plots, 11 differed according to physiographic province, three to calcium levels, and seven to pH levels (Table 5.2).

**Table 5.2.**
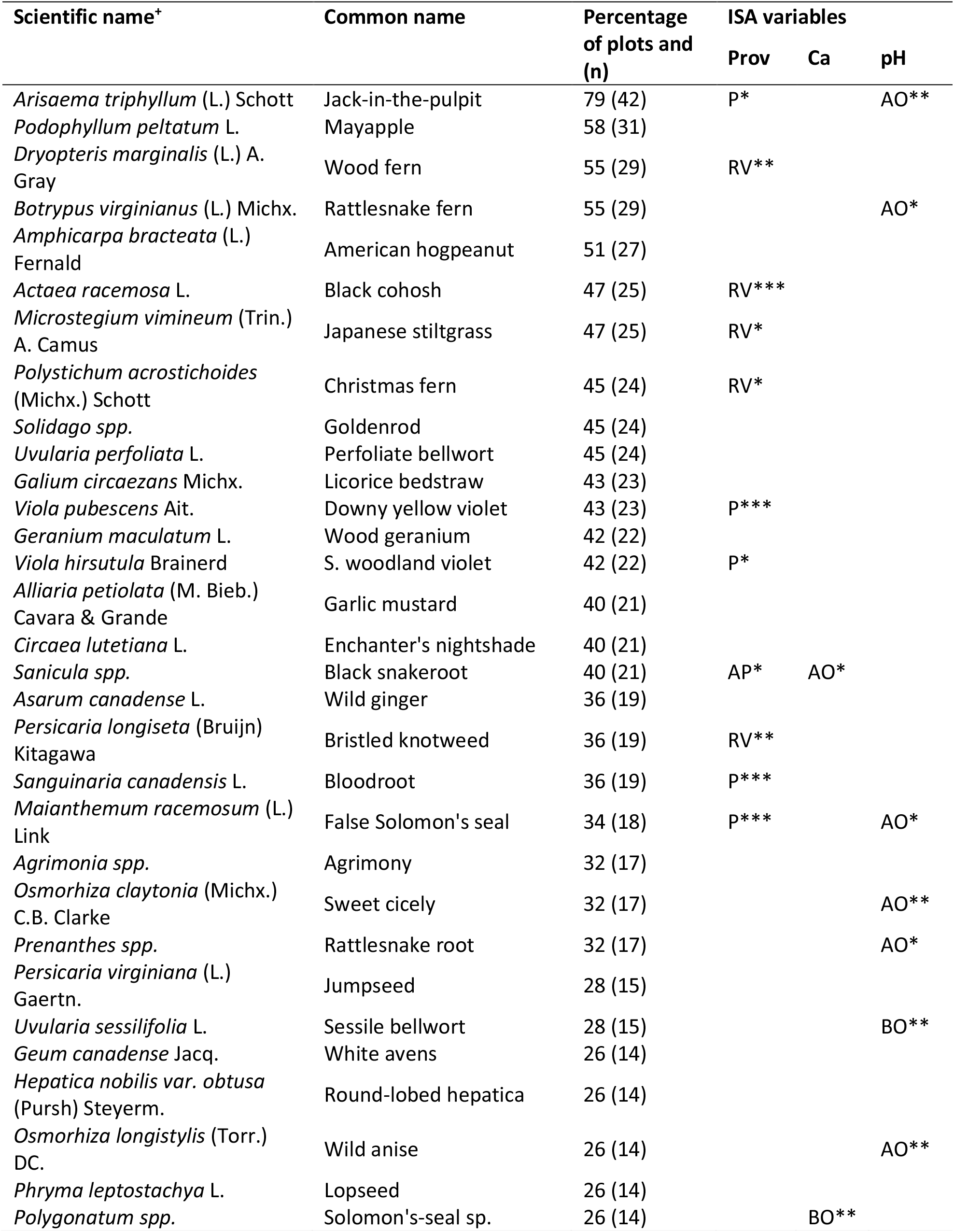
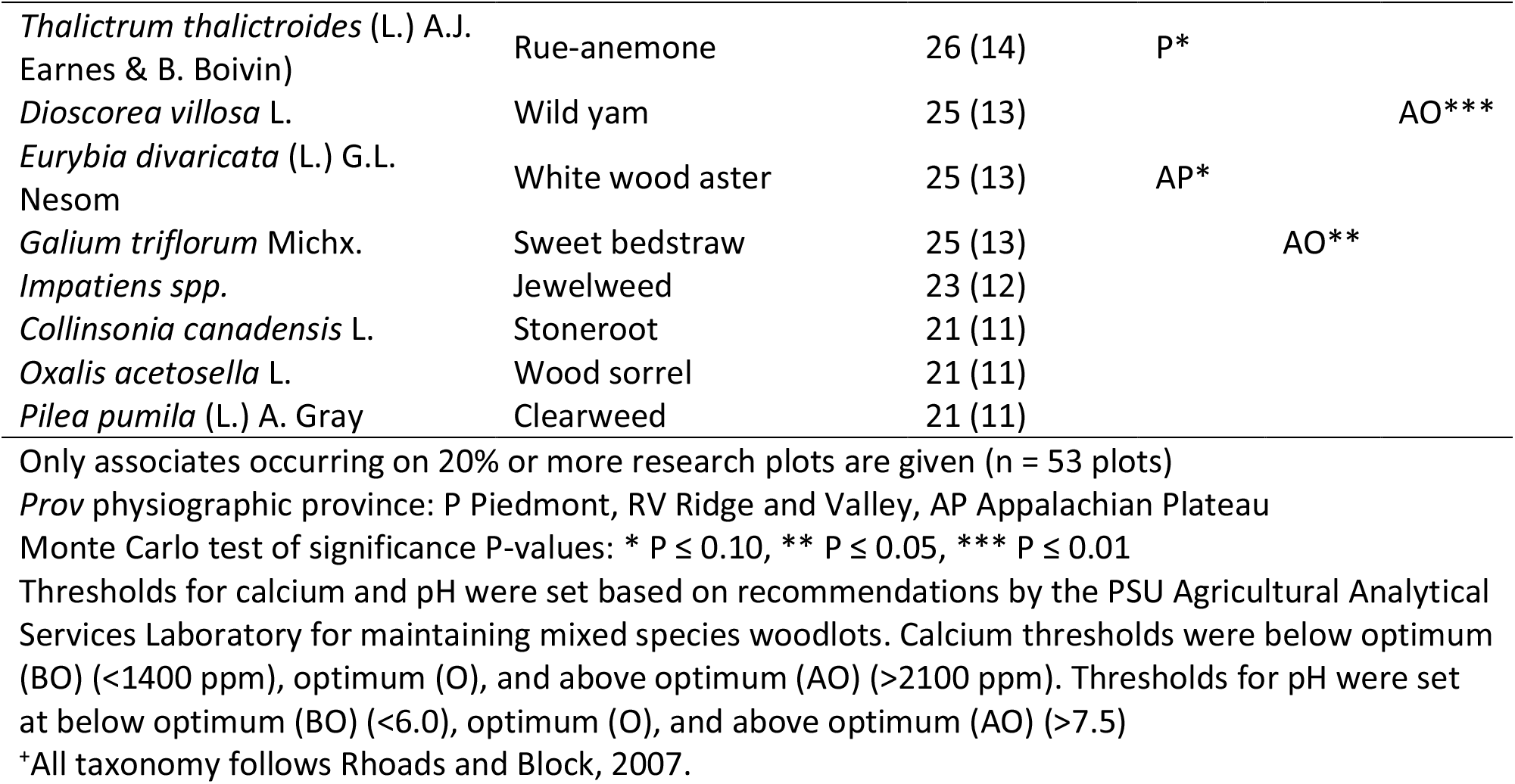
Herbaceous flowering plants and ferns associated with goldenseal in Pennsylvania along with indicator species analysis (ISA) results for physiographic province, calcium, and pH.

Many of the herbaceous species identified in this study have been associates of goldenseal in other states within the species natural range. In Michigan, associated herbs included Jack-in-the-pulpit (*Arisaema triphyllum*), wild ginger *(Asarum canadense*), wild geranium *(Geranium maculatum*), and perfoliate bellwort *(Uvularia perfoliata*), among other herbs commonly found in mesic forests (Penskar et al., 2001). In Virginia, mayapple (*Podophyllum peltatum*), hog peanut (*Amphicarpa bracteata*), black cohosh (*Actaea racemosa*), bloodroot (*Sanguinaria canadensis*), enchanter’s nightshade (*Circaea canadensis*), were identified near a wild goldenseal population (Mueller, 2004).

The most common ferns associated with goldenseal were wood fern (*Dryopteris marginalis* (L.) A. Gray) and rattlesnake fern (*Botrypus virginianus* (L.) Michx.), which were present in 55% of plots (Table 5.2). While previous reports of ferns growing in proximity to goldenseal are sparse, Mueller (2004) identified rattlesnake fern (*Botrypus virginianus*) and Christmas fern (*Polystichum acrostichoides*) growing near a goldenseal population in Virginia. In Pennsylvania, wood ferns (*Dryopteris spp.*) are common in a variety of habitats, including rocky wooded slopes and ravines while rattlesnake fern is commonly found in rich, mesic woods (Rhoads and Block, 2007), and is a common associate of American ginseng as well (Burkhart, 2013).

The most common shrub was spicebush (*Lindera benzoin* L.), which occurred in 83% of the plots. Virginia creeper (*Parthenocissus quinquefolia* (L.) Planch) was the most common vine associate and occurred in 81% of the plots (Table 5.3). Both species have been associated with goldenseal in Virginia (Mueller, 2004).

**Table 5.3.**
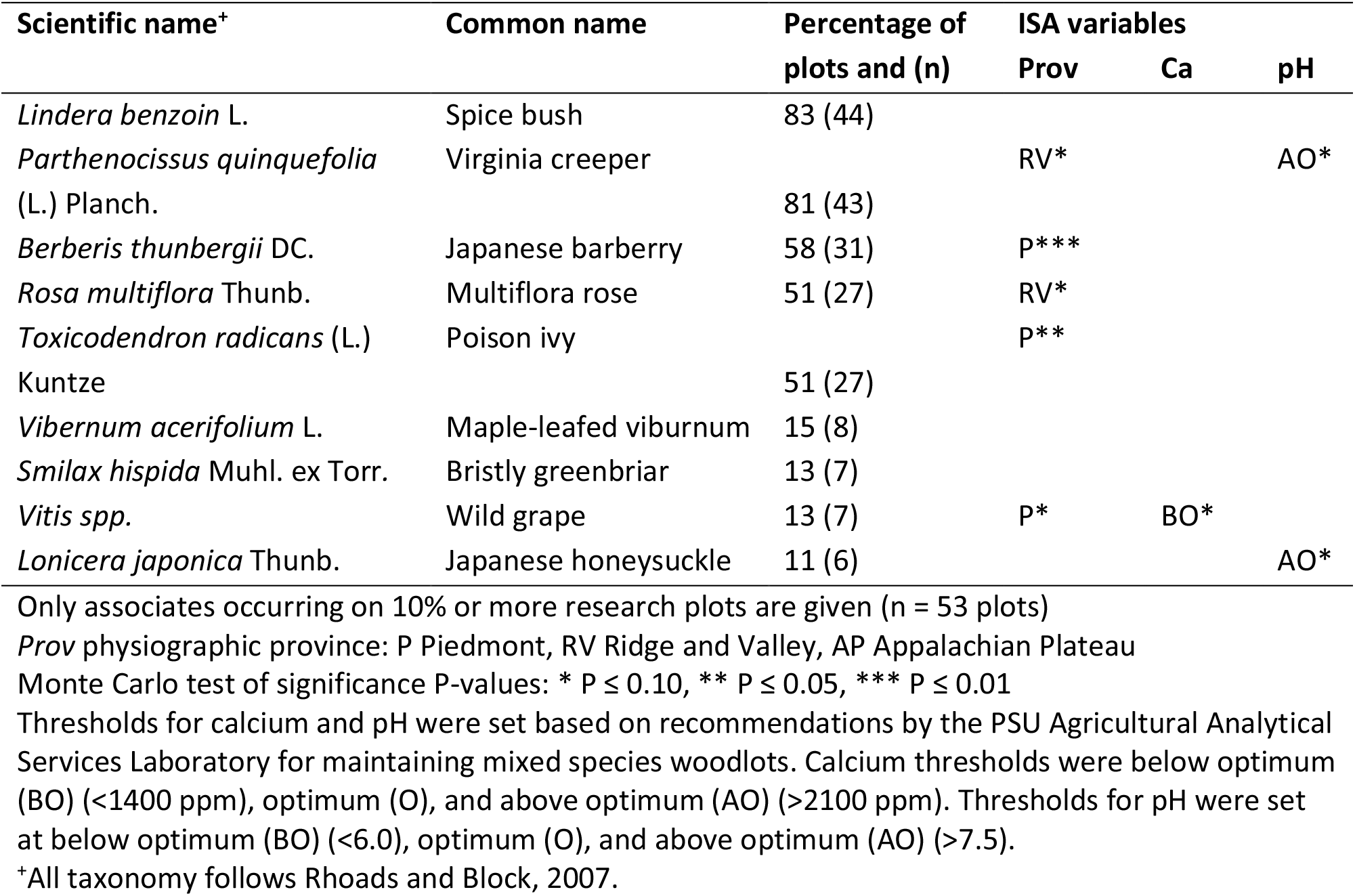
Shrubs and vines associated with goldenseal in Pennsylvania along with indicator species analysis (ISA) results for physiographic province, calcium, and pH.

The most common overstory species were tulip poplar (*Liriodendron tulipifera* L.) and sugar maple (*Acer saccharum* Marshall), which occurred in 40% and 38% of plots respectively (Table 5.4), and are common in mesic woodlands across Pennsylvania (Rhoads and Block, 2007). These species also had the highest importance values of all the associated tree species (Table 5.5). Both of these species, as well as many of the other canopy species identified were also recorded near goldenseal populations in Virginia (Mueller, 2004), and sugar maple was reported near goldenseal in Michigan (Penskar et al., 2001).

**Table 5.4.** Dominant and co-dominant trees associated with goldenseal in Pennsylvania along with indicator species analysis (ISA) results for physiographic province, calcium, and pH.

| Scientific name<sup>+</sup> | Common name | Percentage of plots and (n) | ISA variables |  |  |
| --- | --- | --- | --- | --- | --- |
|  |  |  | Prov | Ca | pH |
| *Liriodendron tulipifera* L. | Tulip-poplar | 40 (23) | P** |  | AO** |
| *Acer saccharum* Marshall | Sugar maple | 38 (22) |  | AO* | O* |
| *Juglans nigra* L. | Black walnut | 22 (13) | RV** |  |  |
| *Carya cordiformis* (Mill.) K. Koch | Bitternut hickory | 16 (9) |  |  | AO*** |
| *Quercus alba* L. | White oak | 16 (9) |  |  |  |
| *Quercus rubra* L. | Northern red oak | 16 (9) |  |  |  |
| *Carya glabra* (Mill.) Sweet | Pignut hickory | 14 (8) | P*** |  | AO*** |
| *Prunus serotina* L. | Black cherry | 12 (7) | AP** |  |  |
| *Tilia americana* L. | American basswood | 10 (6) |  |  |  |
| *Carya tomentosa* (Poir.) Nutt. | Mockernut hickory | 9 (5) | P** | O* | AO** |
| *Carya ovata* (Mill.) K. Koch | Shagbark hickory | 9 (5) |  |  |  |
| *Ulmus rubra* Muhl. | Slippery elm | 9 (5) |  |  |  |
| *Acer platanoides* L. | Norway maple | 5 (3) | AP** |  |  |
| *Fraxinus pennsylvanica* Marshall | Green ash | 5 (3) |  |  | AO* |
| *Quercus montana* Willd. | Chestnut oak | 5 (3) |  |  |  |
| *Fagus grandifolia* Ehrhart | American beech | 5 (3) |  |  |  |

Associates occurring on 5% or more of research plots are given (n = 58 plots)
Prov physiographic province: P Piedmont, RV Ridge and Valley, AP Appalachian Plateau
Monte Carlo test of significance P-values: \* $P \leq 0.10$ , \*\* $P \leq 0.05$ , \*\*\* $P \leq 0.01$
Thresholds for calcium and pH were set based on recommendations by the PSU Agricultural Analytical Services Laboratory for maintaining mixed species woodlots. Calcium thresholds were below optimum (BO) (<1400 ppm), optimum (O), and above optimum (AO) (>2100 ppm). Thresholds for pH were set at below optimum (BO) (<6.0), optimum (O), and above optimum (AO) (>7.5).
<sup>+</sup>All taxonomy follows Rhoads and Block, 2007.

**Table 5.5.**
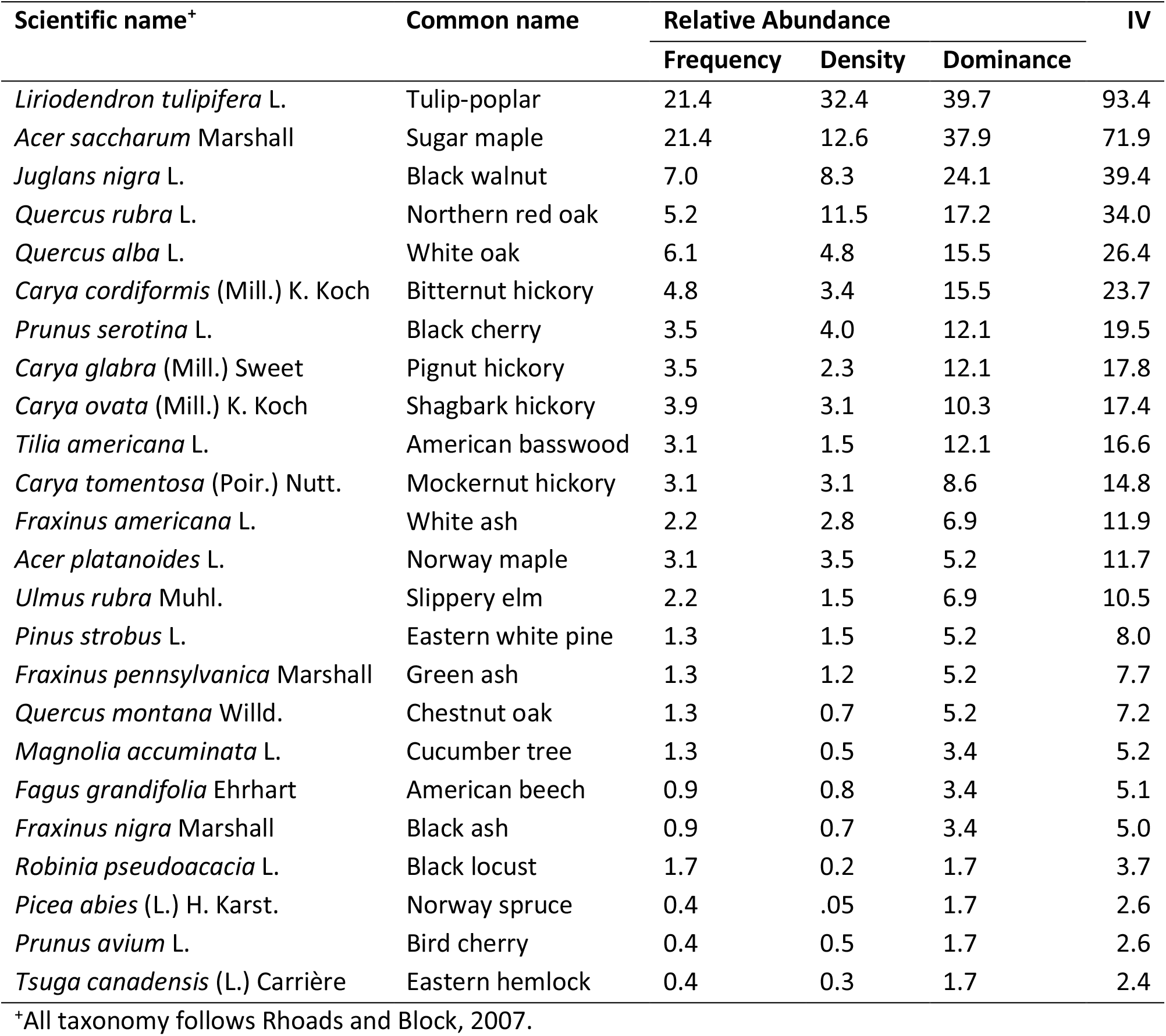
Relative abundances and importance values (IV) for dominant or co-dominant overstory tree species associated with goldenseal in Pennsylvania.

### Site Characteristics

Soil chemistry associated with goldenseal in Pennsylvania varied greatly (Table 5.6). Soil pH ranged from 5 to 7.3, with a mean of 6.2. Macronutrients also varied considerably with standard deviations that were commonly half the value of the mean. Textural class was predominantly loam, with sandy loam, clay loam, and sandy clay loam also recorded. Goldenseal has been documented growing in the same soil textures in Michigan (Penskar et al., 2001).

**Table 5.6.**
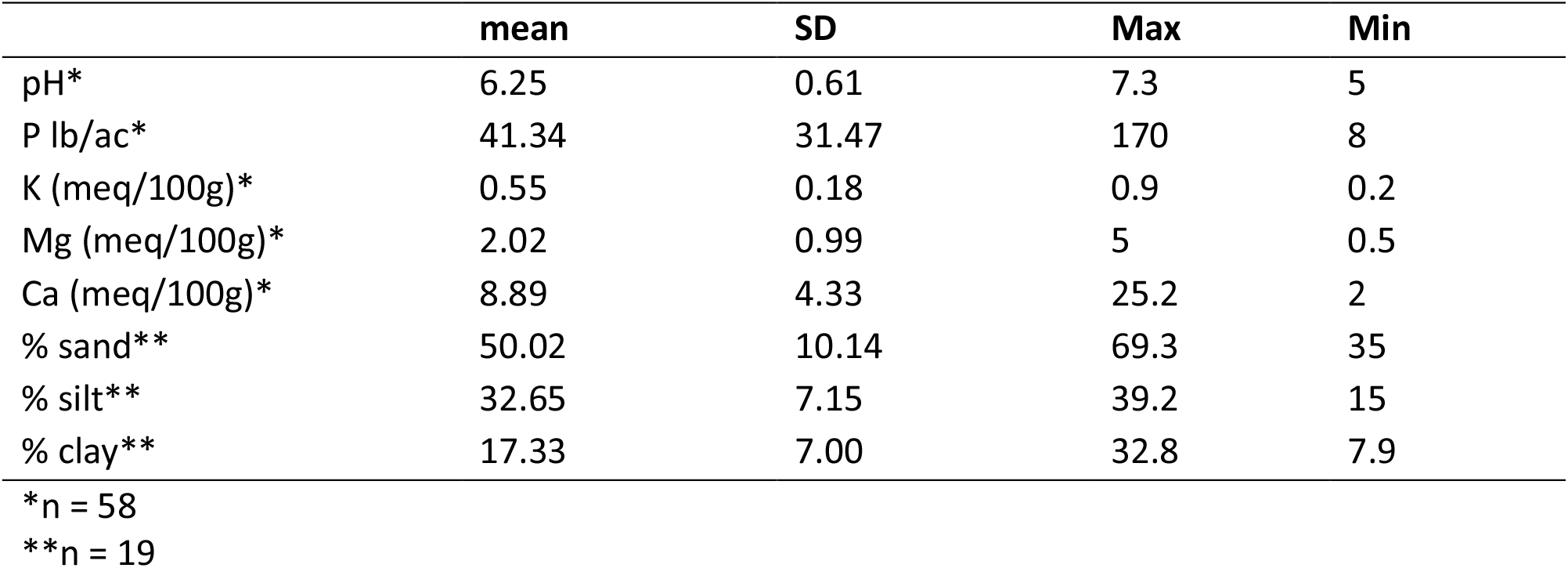
Edaphic factors associated with wild and forest cultivated goldenseal in Pennsylvania.

|  | **mean** | **SD** | **Max** | **Min** |
| --- | --- | --- | --- | --- |
| pH* | 6.25 | 0.61 | 7.3 | 5 |
| P lb/ac* | 41.34 | 31.47 | 170 | 8 |
| K (meq/100g)* | 0.55 | 0.18 | 0.9 | 0.2 |
| Mg (meq/100g)* | 2.02 | 0.99 | 5 | 0.5 |
| Ca (meq/100g)* | 8.89 | 4.33 | 25.2 | 2 |
| % sand** | 50.02 | 10.14 | 69.3 | 35 |
| % silt** | 32.65 | 7.15 | 39.2 | 15 |
| % clay** | 17.33 | 7.00 | 32.8 | 7.9 |

\*n = 58
\*\*n = 19

Elevation at goldenseal sites ranged from 33m to 733m. Populations were found across a range of aspects and topographic positions and from ridge-top to valley bottom, with most of the sites being moist environments (Table B3). The large degree of forested habitat variation across the Pennsylvania populations supports that goldenseal can grow in a variety of site and soil conditions (Upton, 2001; Sinclair and Catling, 2001; Mueller, 2004; Predny and Chamberlain, 2005).

### Maxent Model Performance and Variable Importance

After initial modeling with 50 covariates, the top 10 predictor variables were selected for the final model based on percent contribution threshold of 2%. These variables included three climatic variables, four edaphic variables, and three topographic variables of differing importance (Figure 5.2). The resulting model had a training AUC value of 0.974, and a testing AUC value of 0.991 (Figure 5.3).

**Figure 5.2.**
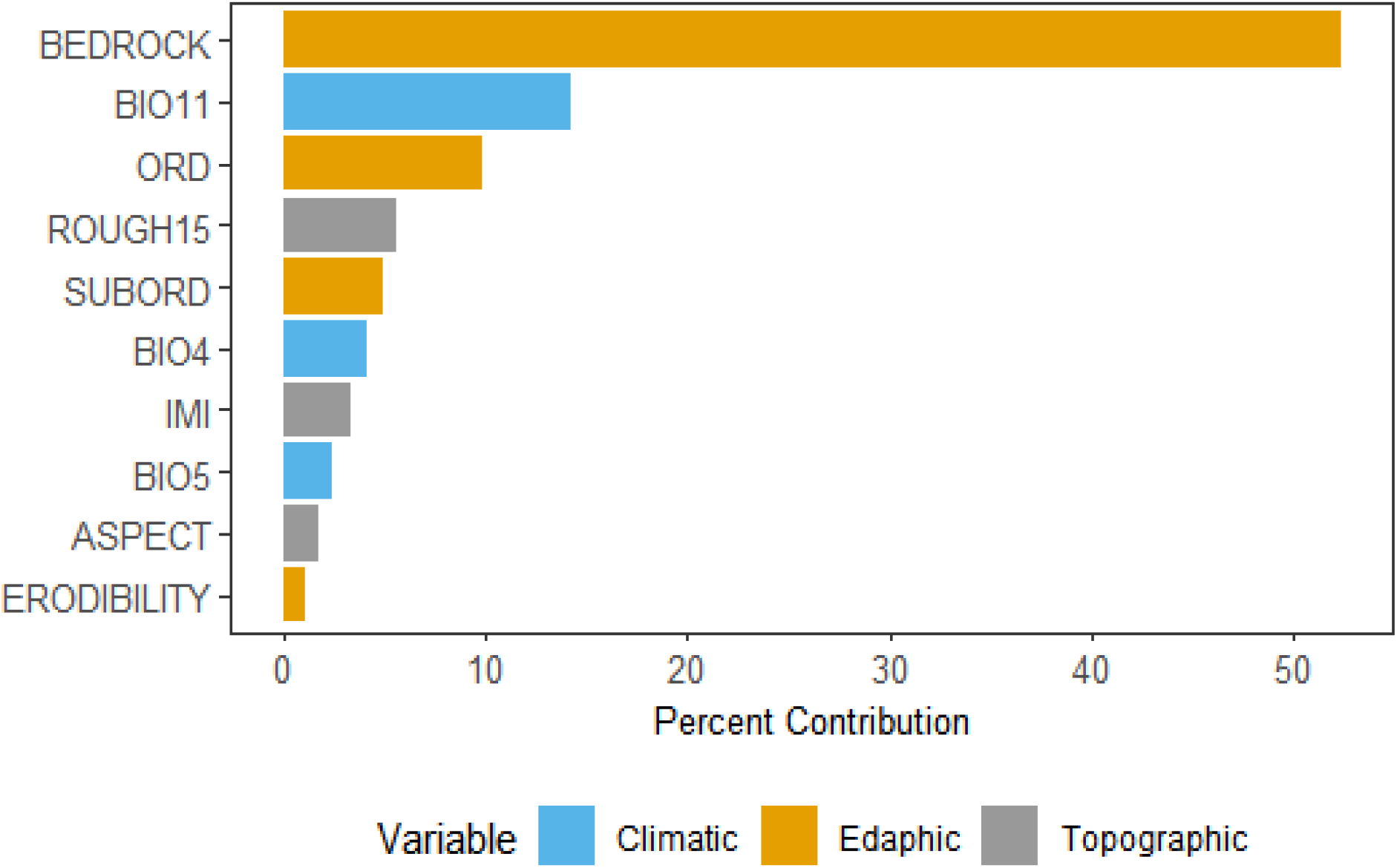
Contribution of the variables used in the final Maxent model.

**Figure 5.3.**
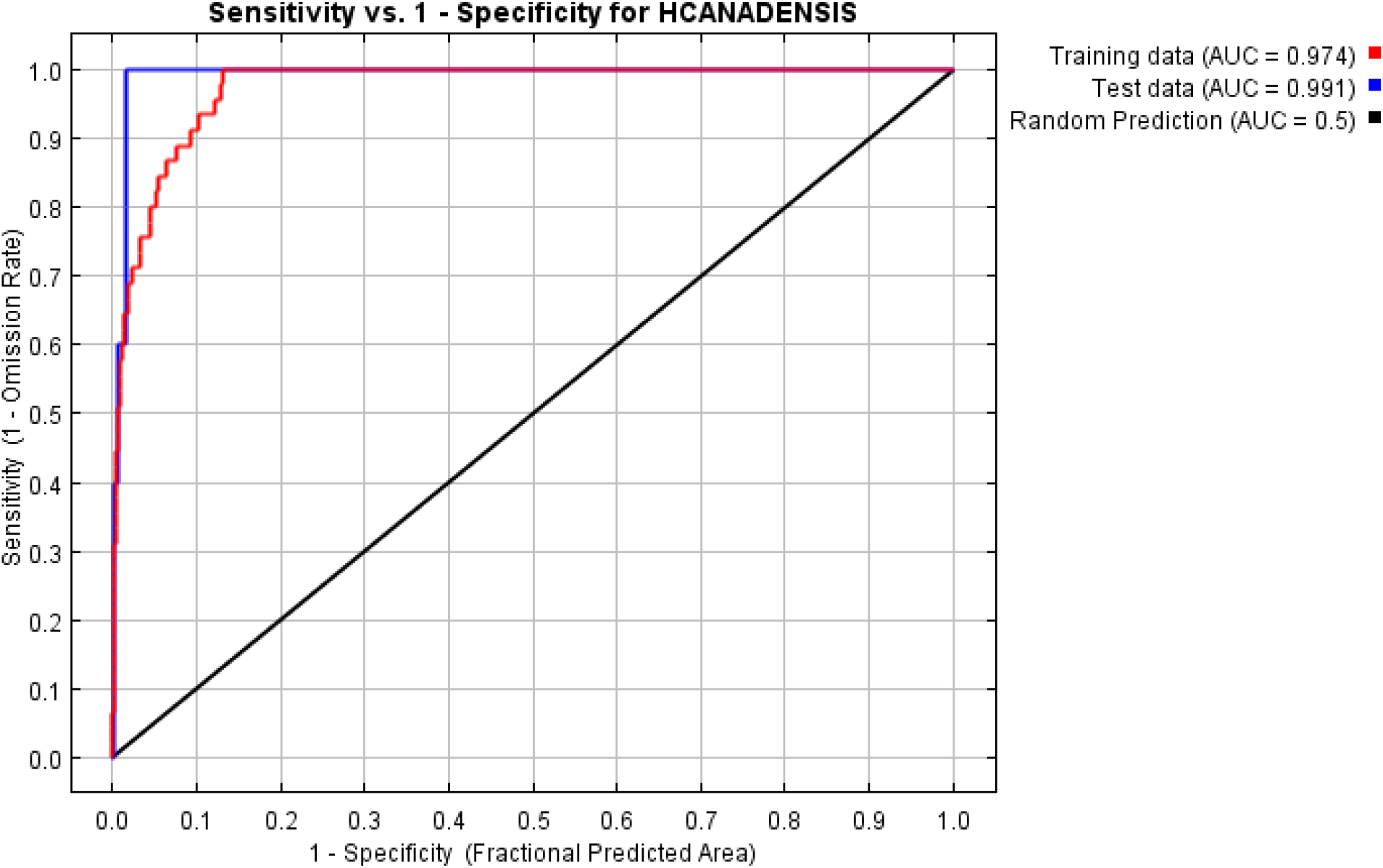
AUC curves from the habitat suitability model for goldenseal. Specificity is established using predicted area, rather than true commission due to having presence-only data.

The two most significant predictor variables as measured by percent contribution and permutation importance were bedrock type and the mean temperature of the coldest quarter (Figure 5.2). Using the jackknife test, results identified bedrock type as the predictor variable that, when isolated, resulted in the greatest gain. Additionally, bedrock type was also identified as the variable that, when removed, resulted in the greatest decrease in gain (Figure 4), indicating that it contains the most information that is not contained in other variables. Response curves of the top ten covariates (Figure 5.5) were used to quantify the suitable habitat for goldenseal (Figure 5.6).

**Figure 5.4.**
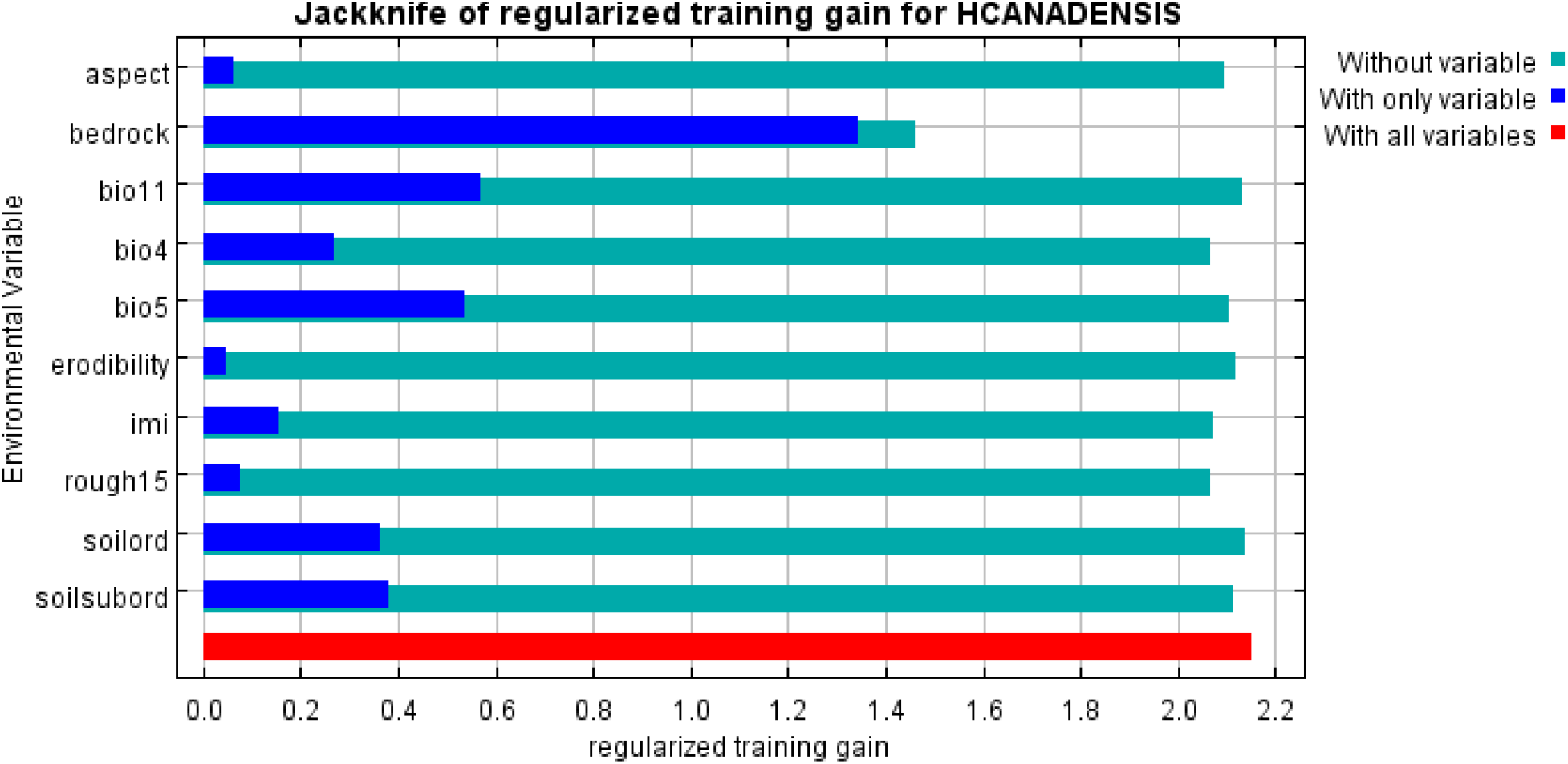
Results of jackknife test of variable importance for modeling goldenseal habitat suitability. The regularized training gain represents how much better the model fits the presence data when compared to a normal distribution. The light green color represents the improvement lost by the removal of that variable from the model, while the dark blue represents the improved fit of the model with that variable alone. The red represents the total improvement from normal of all the variables combined.

**Figure 5.5.**
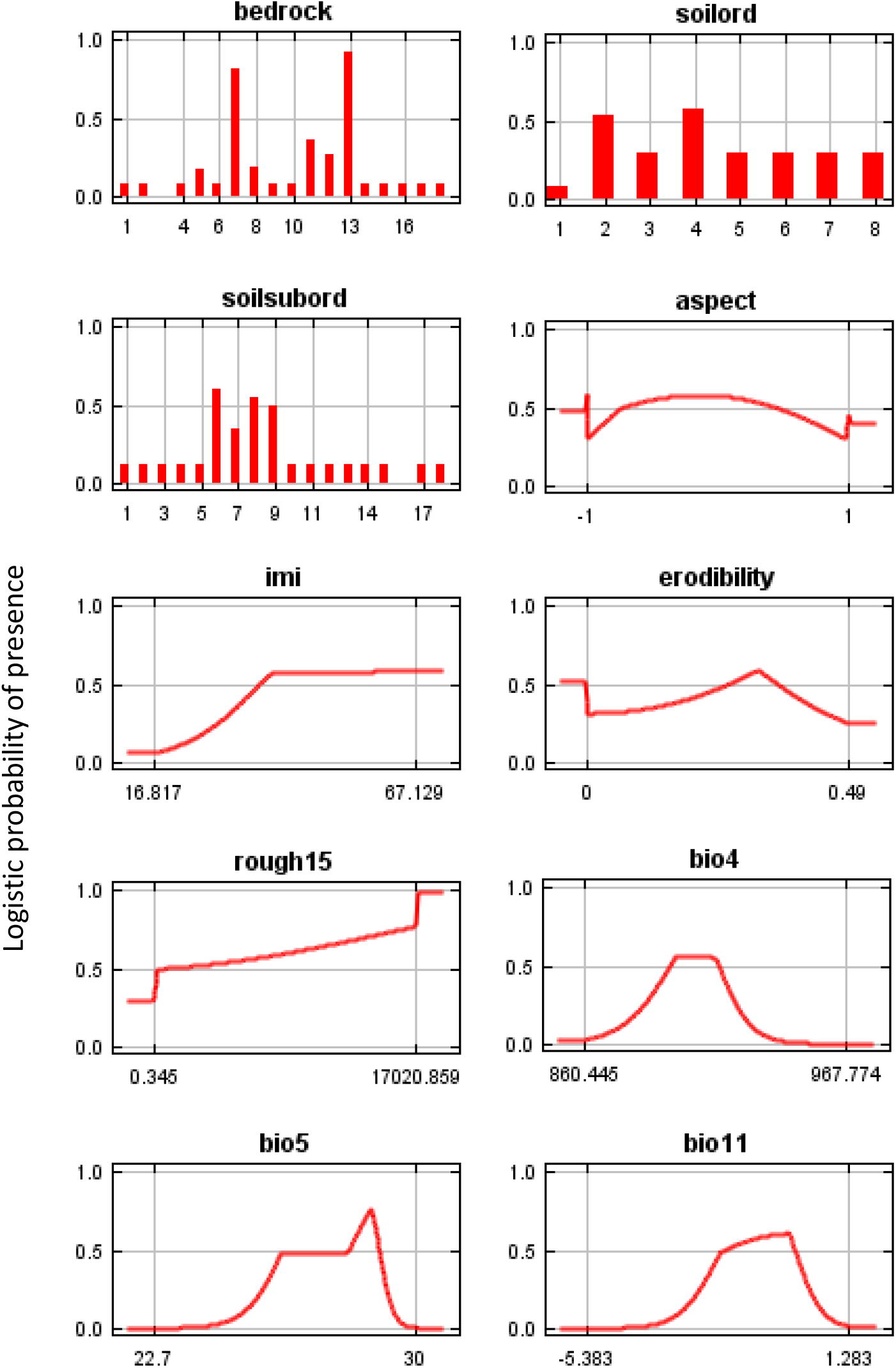
Response curves of the 10 predictor variables included in the final goldenseal habitat distribution model, as created by running the Maxent model using only the corresponding variable. Explanation of each variable can be found in Table 1.

**Figure 5.6.**
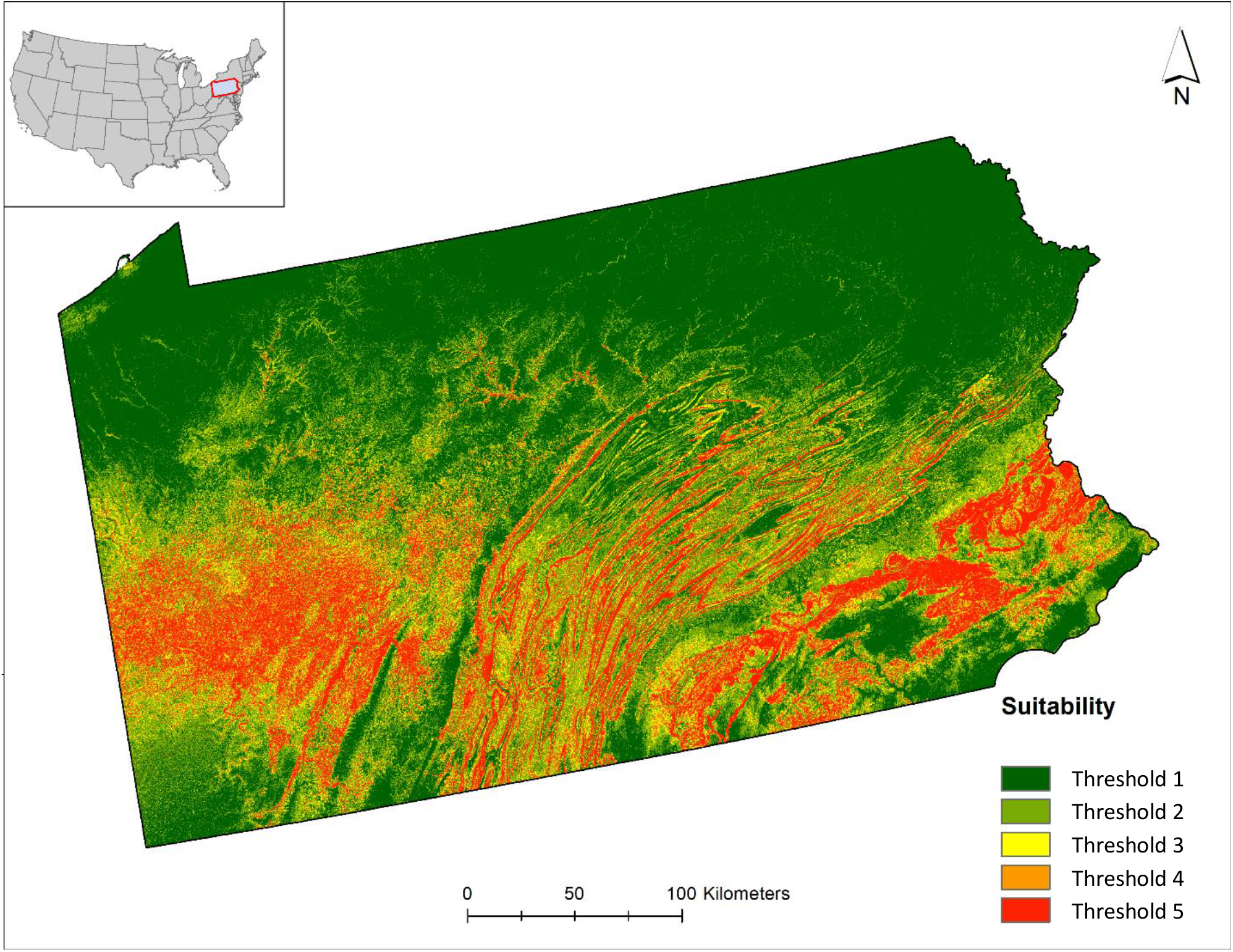
Maxent model for *Hydrastis canadensis* in Pennsylvania. Warmer colors show areas that are predicted to have more suitable habitat conditions. The five thresholds from top to bottom are the fixed cumulative value 1 (roughly 1% of presences would be predicted as absent), fixed cumulative value 5, fixed cumulative value 10, minimum training presence (cumulative threshold of 28), and logistic thresholds for the 10 percentile training presence (cumulative threshold of 34).

### Edaphic Model Results

Bedrock type that increased the suitable habitat for goldenseal was found to be Pennsylvanian in the western part of the state, Devonian and Silurian in the Ridge-and-Valley, and Jurassic in the southeast. Both the Pennsylvanian in the west and the Devonian and Silurian in the Ridge-and-Valley are made up of acidic shale, while the Jurassic in the east is diabase. However, rather than simply indicating ideal habitat for goldenseal, these bedrock types were isolated at least in part because of previous land use. While acidic shale has been farmed in parts of the state, it requires significant soil amendments, and on mountainsides more commonly supports mixed hardwood forests (Blumberg and Cunningham, 1982). The diabase in the east erodes more slowly than the siltstone in the region, resulting in rocky outcroppings that prevent farming (Blumberg and Cunningham, 1982).

The influence of previous land use, what is often termed land use legacy (Bellemare et al., 2002), may be a major factor in restricting goldenseal distribution in fertile limestone soils of the valleys of the Ridge-and-Valley region and piedmont region of Pennsylvania. Mueller (2004) reported goldenseal populations growing on very different bedrock types in Virginia and attributed this to the fact that both sites were not suitable for farming. In Ohio, it was reported that goldenseal was disappearing due to suitable habitat being converted to agricultural uses (Koffler and Gorby, 1957). These rich sites may be well suited for goldenseal where agricultural land has been abandoned; however, outside of the core range, vegetative reproduction is more common than sexual reproduction in goldenseal (Gagnon, 1999; Sanders, 2004), which could be a reason for a lack of recolonizing in reforested fertile sites. In addition to land use legacy, model results indicate some bedrock types may not be suitable for goldenseal. The drier, sandier soils associated with the sandstone ridges of the Ridge-and-Valley region were identified as being less suitable goldenseal habitat in the model

To a lesser degree, soil order was found to influence habitat suitability in the model results. Alfisols, which are common soils under hardwood forests and are known to have a clay-enriched subsoil and high native fertility (Soil Survey Staff, 1999), were most strongly associated with predicted suitable habitat. These findings support previous assertions that goldenseal prefers moist soils, high in organic matter (Penskar et al., 2001; Upton, 2001; Sinclair and Catling, 2001). Two suborders that were found to increase suitability within Alfisols were Aqualfs, which are wet soils, and Udulfs, which are thought to have all been forested at one time. Other suborders that were found to be significant were Aquents, a suborder of Entisols that are usually to permanently wet, and Udults (a suborder of Ultisol). A strong inverse correlation was noted with Inceptisols, which are known for being more freely drained and were negatively associated with predicted habitat.

### Climate Model Results

Mean temperature of the coldest quarter (i.e., the average winter temperature) was the most significant climatic factor in predicting suitable goldenseal habitat. Specifically, once the mean dropped below −2.5°C the probability of suitable habitat quickly declined. Pennsylvania is near the northern edge of goldenseal’s native range, with only a limited number of isolated populations found in states further north and the very southern edge of Ontario, Canada. More commonly, it has been found in the south and west within Pennsylvania, which is contiguous with a proposed core goldenseal range that extends throughout the Ohio river valley (Lloyd and Lloyd, 1884; McGraw et al., 2003; Christensen and Gorchov, 2010; NatureServe, 2018).

In addition to the mean temperature of the coldest quarter, two other climatic variables had a minor influence on the model: temperature seasonality (the amount of temperature variance across a year) and mean maximum temperature of the warmest month. However, the later was likely due to sampling deficiency as no populations were identified along the southern edge of the state, despite the range of goldenseal extending down through southern Appalachia.

### Topographic Model Results

Of the four topographic variables that influenced predicted suitable habitat, the integrated moisture index (IMI) was found to have the greatest influence. IMI is a GIS derived index of soil moisture based on slope, aspect, cumulative flow, curvature of landscape, and the water-holding capacity of the soil (Iverson et al., 1997). In general, the suitability increased with an increase in the IMI, which corroborates previous work that found goldenseal seedling success increase with increased soil moisture (Albrecht and McCarthy, 2006), and supports the assertion that goldenseal prefers mesic conditions (Penskar et al., 2001; Sinclair and Catling, 2001; Rhoads and Block, 2007).

Aspect also had an influence on the model, which has been previously found to significantly influence goldenseal seedling recruitment (Albrecht and McCarthy, 2006). However, it was minimal, which was likely a result of the shortcoming of Beers transformation, which was used for quantifying the aspect in the model. In the transformation, northeast aspect is given the highest value, while southwest is given the lowest. However, the ridges of central Pennsylvania run from southwest to northeast, ultimately resulting in location aspects either being northwest or southeast, which have the same value. The other variables that had a slight influence on the model were an estimate for surface roughness (variance of elevation) and erodibility which together indicated that habitat that is flat is more suited for goldenseal.

## Conclusions

The goal of this study was to identify and quantify existing goldenseal habitat in Pennsylvania through a combination of field collected habitat data and a state-level habitat suitability model. Field studies identified Jack-in-the-pulpit (*Arisaema triphyllum*), wood fern (*Dryopteris marginalis*), rattlesnake fern (*Botrypus virginianus*), and spice bush (*Lindera benzoin*) as the most commonly associated understory species. The most common overstory associates identified were tulip poplar (*Liriodendron tulipifera*) and sugar maple (*Acer saccharum*). Soils were most commonly loams and sandy loams, with and average pH of 6.2 and high variation in macronutrient levels.

The two primary model predictors for goldenseal habitat suitability in PA identified in this study were bedrock type and mean winter temperature. The significance of bedrock type may be a combination of unsuitable habitat and land use legacy. Highly suitable goldenseal habitats were found to be areas where forests were not underlain by sandstone parent materials. In PA, such sites would also be most suited for farming and the occurrence of goldenseal on certain bedrocks may therefore be more a reflection of previous land use rather than any species predilection per se. Average winter temperature was found to be inversely related to habitat suitability. Suitable habitat for goldenseal in Pennsylvania therefore appears to be restricted by the severity of winter temperatures, particularly in the northern part of the state and at higher elevations. This alone may explain why goldenseal is absent from most of northern PA, and has a patchy distribution in northeastern states. It may be that occurrence in the northeast is governed by microsite availability in which winter temperatures are moderated by localized habitat nuances.

When looking for wild populations of goldenseal to conserve, previous land use should be considered in the southern part of the state in which winter temperatures are mild enough to support the species. When looking to reintroduce goldenseal for conservation or forest farming, considering common associates could be an efficient way to identify suitable sites. By utilizing a habitat suitability model combined with associated indicator species analysis, success of conservation, restoration, and reintroduction of goldenseal, and rare species in general, could be increased through more targeted management.

## Appendix

**Table B1.** Total list of variables used in initial Maxent model run

| Variable | Description/Source |
| --- | --- |
| Climatic predictors |  |
| BIO1 | Annual mean temperature (°C) |
| BIO2 | Mean Diurnal Range (Mean of monthly (max temp - min temp)) |
| BIO3 | Isothermality (BIO2/BIO7) (* 100) |
| BIO4 | Temperature Seasonality (standard deviation *100) |
| BIO5 | Max Temperature of Warmest Month |
| BIO6 | Min Temperature of Coldest Month |
| BIO7 | Temperature Annual Range (BIO5-BIO6) |
| BIO8 | Mean Temperature of Wettest Quarter |
| BIO9 | Mean Temperature of Driest Quarter |
| BIO10 | Mean Temperature of Warmest Quarter |
| BIO11 | Mean Temperature of Coldest Quarter |
| BIO12 | Annual Precipitation |
| BIO13 | Precipitation of Wettest Month |
| BIO14 | Precipitation of Driest Month |
| BIO15 | Precipitation Seasonality (Coefficient of Variation) |
| BIO16 | Precipitation of Wettest Quarter |
| BIO17 | Precipitation of Driest Quarter |
| BIO18 | Precipitation of Warmest Quarter |
| BIO19 | Precipitation of Coldest Quarter |
| FFP | Length of the frost-free period |
| Edaphic predictors |  |
| AWC | Available water capacity (cm) |
| BEDROCK | Bedrock type |
| BD | Soil bulk density (g/cm<sup>3</sup>) |
| DEPTH | Depth to bedrock (cm) |
| ERODIBILITY | Soil erodibility factor, rock fragment free |
| OM | Organic matter content (% by weight) |
| CLAY | Percent clay |
| SAND | Percent sand |
| SILT | Percent silt |
| PERM | Soil permeability rate (cm/hr) |
| PH | Soil pH |
| NO10 | Percent soil passing sieve no.10 (coarse) |
| NO200 | Percent soil passing sieve no.200 (fine) |
| SOILSP | Soil slope(%) of a soil component |
| ORD | Soil order based on USDA soil taxonomy |
| SUBORD | Soil suborder based on USDA soil taxonomy |
|  | Topographic predictors |
| AP | Transformed aspect according to Beer et al. (1966) |
| ERR15 | Elevation relief ratio (15-pixel neighborhood) |
| IMI | Integrated soil moisture index (Iverson et al. 1997) |
| ROUGH15, 27 | Variance of elevation (15-pixel, 27-pixel neighborhood) |
| SP | Slope in radians |
| SPCOSAP | Slope x COS(Aspect) |
| SPSINAP | Slope x SIN(Aspect) |
| TRASP | Topographic radiation index |
| TPI150, 300, 500,<br>1000, 2000 | Topographic position indices at scales of 150m, 300m, 500m,<br>1000m, and 2000m |

**Table B2.** Frequency of species identified as being associated with goldenseal in Pennsylvania (n = 53 plots)

| Scientific name* | Common name | Percentage of plots | Number of plots |
| --- | --- | --- | --- |
| *Lindera benzoin* (L.) Blume | Spicebush | 83 | 44 |
| *Parthenocissus quinquefolia* (L.) Planch | Virginia creeper | 81 | 43 |
| *Arisaema triphyllum* (L.) Schott | Jack-in-the-pulpit | 79 | 42 |
| *Berberis thunbergii* DC | Japanese barberry | 58 | 31 |
| *Podophyllum peltatum* L. | Mayapple | 58 | 31 |
| *Botrypus virginianus* (L.) Michx. | Rattlesnake fern | 55 | 29 |
| *Dryopteris marginalis* (L.) A. Gray | Marginal shield fern | 55 | 29 |
| *Amphicarpa bracteata* (L.) Fernald | American hogpeanut | 51 | 27 |
| *Rosa multiflora* Thumb. ex Murray | Multiflora rose | 51 | 27 |
| *Toxicodendron radicans* (L.) Kuntze | Poison ivy | 51 | 27 |
| *Actaea racemosa* L. | Black cohosh | 47 | 25 |
| *Microstegium vimineum* (Trin.) A. Camus | Japanese stiltgrass | 47 | 25 |
| *Polystichum acrostichoides* (Michx.) Schott | Christmas fern | 45 | 24 |
| *Uvularia perfoliata* L. | Perfoliate bellwort | 45 | 24 |
| *Galium circaeazans* Michx. | Licorice bedstraw | 43 | 23 |
| *Liriodendron tulipifera* L. | Tulip-poplar | 43 | 23 |
| *Viola pubescens* Ait. | Downy yellow violet | 43 | 23 |
| *Acer saccharum* Marshall | Sugar maple | 42 | 22 |
| *Geranium maculatum* L. | Wood/wild geranium | 42 | 22 |
| *Viola hirsutula* Brainerd | Southern woodland violet | 42 | 22 |
| *Alliaria petiolata* (M. Bieb.) Cavara & Grande | Garlic mustard | 40 | 21 |
| *Circaea canadensis* L. | Enchanter's nightshade | 40 | 21 |
| *Sanicula* spp. | Black snakeroot | 40 | 21 |
| *Asarum canadense* L. | Wild ginger | 36 | 19 |
| *Persicaria longiseta* (Bruijn) Kitagawa | Bristled knotweed | 36 | 19 |
| *Sanguinaria canadensis* L. | Bloodroot | 36 | 19 |
| *Maianthemum racemosum* (L.) Link | False Solomon's seal | 34 | 18 |
| *Agrimonia* spp. | Agrimony | 32 | 17 |
| *Osmorhiza claytonii* (Michx.) C. B. Clarke | Clayton's sweetroot | 32 | 17 |
| *Prenanthes* spp. | Rattlesnake root | 32 | 17 |
| *Persicaria virginiana* (L.) Gaertn. | Virginia knotweed | 28 | 15 |
| *Uvularia sessilifolia* L. | Sessile bellwort | 28 | 15 |
| *Geum canadense* Jacq. | White avens | 26 | 14 |
| *Hepatica nobilis* var. *obtusata* (Pursh) Steyerl. | Round-lobed hepatica | 26 | 14 |
| *Osmorhiza longistylis* (Torr.) DC. | Wild anise | 26 | 14 |
| *Phryma leptostachya* L. | Lopseed | 26 | 14 |
| *Thalictrum thalictroides* (L.) A.J. Earnes & B. Boivin | Rue-anemone | 26 | 14 |
| *Dioscorea villosa* L. | Wild yam | 25 | 13 |
| *Eurybia divaricata* (L.) G.L. Nesom | White wood aster | 25 | 13 |
| *Galium triflorum* Michx. | Sweet bedstraw | 25 | 13 |
| *Juglans nigra* L. | Black Walnut | 25 | 13 |
| *Collinsonia canadensis* L. | Canada horsebalm | 21 | 11 |
| *Oxalis acetosella* L. | Wood sorrel | 21 | 11 |
| *Pilea pumila* (L.) A. Gray | Clearweed | 21 | 11 |
| *Galearis spectabilis* (L.) Raf. | Showy orchid | 19 | 10 |
| *Ageratina altissima* (L.) King & H.E. Robins. | White snakeroot | 17 | 9 |
| *Cardamine concatenata* (Michx.) O. Schwarz | Cutleaf toothwort | 17 | 9 |
| *Quercus alba* L. | White oak | 17 | 9 |
| *Quercus rubra* L. | Northern red oak | 17 | 9 |
| *Carya glabra* (Mill.) Sweet | Pignut hickory | 15 | 8 |
| *Erythronium americanum* Ker-Gawl. | Yellow trout-lily | 15 | 8 |
| *Hackelia virginiana* I.M. Johnston | Beggar's lice | 15 | 8 |
| *Impatiens capensis* Meerb. | Common jewelweed | 15 | 8 |
| *Viburnum acerifolium* L. | Maple-leaved viburnum | 15 | 8 |
| *Adiantum pedatum* L. | Maidenhair fern | 13 | 7 |
| *Claytonia virginica* L. | Virginia springbeauty | 13 | 7 |
| *Monarda didyma* L. | Scarlet beebalm | 13 | 7 |
| *Monotropa uniflora* L. | Indian pipe | 13 | 7 |
| *Panax quinquefolius* L. | American ginseng | 13 | 7 |
| *Prosartes lanuginosa* (Michx.) D. Don | Yellow mandarin | 13 | 7 |
| *Prunus serotina* L. | Black Cherry | 13 | 7 |
| *Smilax hispida* Muhl. ex Torr. | Bristly greenbriar | 13 | 7 |
| *Vitis* spp. | Wild grape | 13 | 7 |
| *Cryptotaenia canadensis* (L.) DC. | Canadian honewort | 11 | 6 |
| *Lonicera japonica* Thunb. | Japanese honeysuckle | 11 | 6 |
| *Polygonatum biflorum* (Walt.) Ell. | Smooth Solomon's-seal | 11 | 6 |
| *Smilax herbacea* L. | Smooth carrionflower | 11 | 6 |
| *Tilia americana* L. | American basswood | 11 | 6 |
| *Allium tricoccum* Aiton | Ramps | 9 | 5 |
| *Carya ovata* (Mill.) K. Koch | Shagbark hickory | 9 | 5 |
| *Carya tomentosa* (Poir.) Nutt. | Mockernut hickory | 9 | 5 |
| *Fragaria vesca* L. | Wild strawberry | 9 | 5 |
| *Maianthemum canadense* Desf. | Canada mayflower | 9 | 5 |
| *Polygonatum pubescens* (Willd.) Pursh | Hairy Solomon's-seal | 9 | 5 |
| *Trillium erectum* L. | Purple trillium/bethroot | 9 | 5 |
| *Ulmus rubra* Muhl. | Red elm | 9 | 5 |
| *Carya cordiformis* (Wang) K. Koch | Bitternut hickory | 8 | 4 |
| *Celastrus orbiculatus* Thunb. | Oriental bittersweet | 8 | 4 |
| *Desmodium* spp. | Tick trefoil | 8 | 4 |
| *Mitchella repens* L. | Partridge berry | 8 | 4 |
| *Onoclea sensibilis* L. | Sensitive fern | 8 | 4 |
| *Rubus phoenicolasius* Maxim | Wineberry | 8 | 4 |
| *Sedum ternatum* Michx. | Woodland stonecrop | 8 | 4 |
| *Smilax rotundifolia* L. | Greenbriar | 8 | 4 |
| *Thelypteris noveboracensis* (L.) Nieuwl. | New York Fern | 8 | 4 |
| *Actaea pachypoda* Elliott | White baneberry/doll's-eyes | 6 | 3 |
| *Acer platanoides* L. | Norway maple | 6 | 3 |
| *Anemone quinquefolia* L. | Wood anemone | 6 | 3 |
| *Caulophyllum thalictroides* L. | Blue cohosh | 6 | 3 |
| *Clinopodium vulgare* L. | Wild basil | 6 | 3 |
| *Euonymus alatus* (Thunb.) Siebold | Winged euonymous | 6 | 3 |
| *Fagus grandifolia* Ehrhart | American beech | 6 | 3 |
| *Fraxinus pennsylvanica* Marshall | Green ash | 6 | 3 |
| *Laportea canadensis* (L.) Wedd. | Wood-nettle | 6 | 3 |
| *Phegopteris hexagonoptera* (Michx.) Fée | Beech fern | 6 | 3 |
| *Quercus montana* Willd. | Chestnut oak | 6 | 3 |
| *Solidago flexicaulis* L. | Broadleaved goldenrod | 6 | 3 |
| *Trillium grandiflorum* (Michx.) Salisb. | Large-flowered trillium | 6 | 3 |
| *Tussilago farfara* L. | Coltsfoot | 6 | 3 |
| *Aralia racemosa* L. | American spikenard | 4 | 2 |
| *Artemisia* spp. | Mugwort | 4 | 2 |
| *Erechtites hieraciifolius* (L.) Raf. Ex DC. | Pilewort | 4 | 2 |
| *Eutrochium* spp. | Joe-Pye weed | 4 | 2 |
| *Fraxinus americana* L. | White ash | 4 | 2 |
| *Fraxinus nigra* Marshall | Black ash | 4 | 2 |
| *Hamamelis virginiana* L. | Witch-hazel | 4 | 2 |
| *Helianthus* spp. | Sunflower | 4 | 2 |
| *Magnolia accuminata* L. | Cucumber tree | 4 | 2 |
| *Packera aurea* (L.) W.A. Weber & Á. Löve | Golden ragwort | 4 | 2 |
| *Pinus strobus* L. | Eastern white pine | 4 | 2 |
| *Ranunculus abortivus* L. | Littleleaf buttercup | 4 | 2 |
| *Sassafras albidum* (Nutt.) Nees | Sassafras | 4 | 2 |
| *Tiarella cordifolia* L. | Heartleaf foamflower | 4 | 2 |
| *Viburnum prunifolium* L. | Blackhaw | 4 | 2 |
| *Viola rostrata* Pursh | Longspur violet | 4 | 2 |
| *Viola striata* Aiton | Striped violet | 4 | 2 |
| *Acer pennsylvanicum* L. | Striped maple | 2 | 1 |
| *Anemone canadensis* L. | Canada anemone | 2 | 1 |
| *Anemone virginiana* L. | Tall thimbleweed | 2 | 1 |
| *Aplectrum hyemale* (Muhl. Ex Willd.) Butt. | Putty-root orchid | 2 | 1 |
| *Blephilia ciliata* (L.) Benth. | Downy wood mint | 2 | 1 |
| *Campanula americana* L. | Tall bellflower | 2 | 1 |
| *Clintonia umbellulata* (Michx.) Morong | White clintonia | 2 | 1 |
| *Cretagus spp.* | Hawthorn | 2 | 1 |
| *Cynoglossum virginianum* L. | Wild comfrey | 2 | 1 |
| *Daucus carota* L. | Wild carrot | 2 | 1 |
| *Desmodium nudiflorum* (L.) DC. | Naked-flowered tick trefoil | 2 | 1 |
| *Dicentra cucullaria* (L.) Bernh. | Dutchman's breeches | 2 | 1 |
| *Eupatorium altissimum* L. | Tall boneset | 2 | 1 |
| *Fallopia* sp. | Bindweed | 2 | 1 |
| *Galium aparine* L. | Sticky willy | 2 | 1 |
| *Galium lanceolatum* (Torr. & A. Gray) Torr. | Lanceleaf wild licorice | 2 | 1 |
| *Hesperis matronalis* L. | Dames rocket | 2 | 1 |
| *Hydrophyllum virginianum* L. | Virginia waterleaf | 2 | 1 |
| *Impatiens pallida* Nutt. | Yellow jewelweed | 2 | 1 |
| *Lysimachia ciliata* L. | Fringed loosestrife | 2 | 1 |
| *Menispermum canadense* L. | Common moonseed | 2 | 1 |
| *Medeola virginiana* L. | Indian cucumber-root | 2 | 1 |
| *Obolaria virginica* L. | Virginia pennywort | 2 | 1 |
| *Persicaria perfoliata* (L.) H. Gross | Mile-a-minute weed | 2 | 1 |
| *Phytolacca americana* L. | American pokeweed | 2 | 1 |
| *Phlox divaricate* L. | Woodland phlox | 2 | 1 |
| *Picea abies* (L.) H. Karst. | Norway spruce | 2 | 1 |
| *Plantago* sp. | Plantain | 2 | 1 |
| *Prunus avium* L. | Bird cherry | 2 | 1 |
| *Prunella vulgaris* L. | Self-heal | 2 | 1 |
| *Robinia pseudoacacia* L. | Black locust | 2 | 1 |
| *Rubus occidentalis* L. | Black raspberry | 2 | 1 |
| *Staphylea trifolia* L. | American bladdernut | 2 | 1 |
| *Symplocarpus foetidus* Salisb. | Skunk cabbage | 2 | 1 |
| *Teucrium canadense* L. | American germander | 2 | 1 |
| *Thalictrum dioicum* L. | Early meadow-rue | 2 | 1 |
| *Tsuga canadensis* (L.) Carrière | Eastern hemlock | 2 | 1 |
| *Verbena urtisiifolia* L. | Nettle-leaved vervain | 2 | 1 |
| *Viola blanda* Willd. | Sweet white violet | 2 | 1 |
| *Viola canadensis* L. | Canadian white violet | 2 | 1 |

\*All taxonomy follows Rhoads and Block, 2007.

**Table B3.** Study sites, population traits and habitat characteristics associated with goldenseal in Pennsylvania

| Site Name | Population size and area | Number of sample plots | Site description | Aspect | Elevation (M) | Topographic position |
| --- | --- | --- | --- | --- | --- | --- |
| Allegheny-SH | 500 ramets, 20% reproductive, <1 hectare | 3 | Moist, lower-slope | W, SW | 251 | Lower-slope |
| Beaver-CM | 1,000 ramets, 20% reproductive, >1 hectare | 2 | Moist side-slope bench | N, NW | 235 | Upper-slope |
| Bedford-BM | 5,000 ramets, 5% reproductive, >1 hectare | 3 | Mid-slope, recent logging | W | 423 | Mid-slope |
| Bedford-WW | >10,000 ramets, 15% reproductive, >1 hectare | 4 | Moist, forest farmed site | None | 348 | Flat |
| Bucks-QT | 5,000-10,000 ramets, 10% reproductive >1 hectare | 4 | moist lowland | None | 119 | Flat |
| Butler-JE | 100-500 ramets, 30% reproductive, <1 hectare | 1 | Moist lower-slope | W | 341 | Lower-slope |
| Cambria-GL | 100-500 ramets, <5% reproductive | 1 | Moist ridgetop | none | 733 | ridgetop |
| Cambria-JK | 500-1,000 ramets, 25% reproductive <1 hectare | 1 | Upland, forest farmed | none | 521 | Top of a hill |
| Centre-ST | >10,000 ramets, 5% reproductive, >1 hectare | 4 | Moist side-slope on drainage bench | NW | 336 | Mid-slope |
| Chester-LF | >10,000 ramets, 5% reproductive, >1 hectare | 4 | Moist lower-slope to dry ridge | N, NE | 104 | Lower-slope, mid-slope, upper-slope |
| Clarion-LH | 50-100 ramets, <5% reproductive, <1 hectare | 1 | Mid-slope in steep ravine | E | 319 | Mid-slope |
| Daughin-JM | 50-100 ramets, 60% reproductive, <1 hectare | 1 | Moist lowland | none | 213 | flat |
| Huntingdon-MG | 5,000 ramets, 15% reproductive, >1 hectare | 4 | Gently sloping mountain hollow | none | 232 | Hollow bottom |
| Indiana-PR | 5,000-10,000 ramets, <5% reproductive, >1 hectare | 4 | Moist, toe-slope, lowland | N | 357 | Lower-slope, valley bottom |
| Lancaster-BW | 1,000 ramets, 5% reproductive, >1 hectare | 4 | Moist lowland | None | 115 | Flat |
| Lancaster-MV | 1,000-5,000 ramets, 15% reproductive, <1 hectare | 1 | Moist lowland | none | 176 | flat |
| Lebanon-GD | 10-50 ramets, <5% reproductive, <1 hectare | 1 | Moist flat site | none | 201 | flat |
| Lebanon-GL | 100 ramets,<br>5%<br>reproductive,<br><1 hectare | 1 | Wet<br>lowland | none | 188 | flat |
| Mifflin-JB | 1,000 ramets,<br>10%<br>reproductive,<br><1 hectare | 3 | Moist<br>sinkhole | SE | 196 | Lower-slope |
| Montgomery<br>-BL | 1000-5000<br>ramets, <5%<br>reproductive,<br>1 hectare | 1 | Moist<br>lower-slope | N | 40 | Lower-slope |
| Montgomery<br>-CD | 100 ramets,<br><5%<br>reproductive,<br><1 hectare | 1 | Moist<br>lowland | none | 57 | flat |
| Montgomery<br>-GL | 100-500<br>ramets, 15%<br>reproductive,<br>1 hectare | 1 | Moist mid-slope | NE | 58 | Mid-slope |
| Montgomery<br>-SX | 100-500<br>ramets, 10%<br>reproductive,<br><1 hectare | 1 | Moist,<br>mid-slope<br>bench | NW | 33 | Mid-slope |
| Westmoreland-DP | 100-500<br>ramets, 10%<br>reproductive,<br><1 hectare | 1 | Moist mid-slope | N | 329 | Mid-slope |
| Westmoreland-MA | >10,000<br>ramets, 5%<br>reproductive,<br>1 hectare | 2 | Moist, flat<br>site | none | 287 | flat |
| Westmoreland-PM | 500-1,000 ramets, 15% reproductive, <1 hectare | 1 | Wet lowland site | none | 377 | flat |
| York-AB | 1,000 ramets, 5% reproductive, 1 hectare | 1 | Wet lowland | none | 183 | flat |
| York-GP | 5,000 ramets, 5% reproductive, >1 hectare | 3 | Dry side-slope and moist bench | SE, NW | 133 | Mid-slope |

